# Decentralizing genetic testing for biodiversity monitoring and biosurveillance with the Nucleic Acid Barcode Identification Tool (NABIT) and Molecular Development Kit (MDK)

**DOI:** 10.1101/2024.03.01.582993

**Authors:** HR Holmes, M Winters, C Fang, G Fotouhi, J Mercader, DA Fox, PME Bunje, A Dehgan

## Abstract

1. The escalating threats to biodiversity, public health, and food security posed by emerging infectious diseases and illegal wildlife trafficking requires novel approaches to biosurveillance. This paper introduces two innovations developed to address these multifaceted challenges: the Nucleic Acid Barcode Identification Tool (NABIT) and the Molecular Development Kit (MDK).
2. The NABIT is a handheld, battery-powered device that enables genetic tests to be performed at the point of contact by non-technical users, creating a critical bridge between centralized laboratories and the field by reducing barriers to accessible and routine genetic testing. Verification testing and validation results for the NABIT and the lyophilized assays used with it demonstrate key performance parameters, including sample preparation, detection sensitivity, and stability of field-ready assays after 17 months without refrigeration.
3. The MDK complements the NABIT by providing a framework for third-party development and deployment of field-ready genetic assays. Similar to software development kits (SDKs), the MDK offers documentation, software tools, and NABIT hardware to accelerate the development of new assays, enabling early detection strategies and minimizing future losses. Crucially, the MDK empowers scientists worldwide to contribute to a new ecosystem for wildlife genetics and biosurveillance by developing their own field-ready tests for the NABIT.
4. In summary, the NABIT and MDK present the potential for a paradigm shift in biosurveillance, ecosystem monitoring, and biodiversity conservation, enabling decentralized genetic testing, early disease detection, and rapid response to protect sensitive ecosystems, public health, and food security.

## 1. Introduction

The IUCN lists over 42,000 species as under threat, and over 16,000 species believed to be endangered for extinction.^1^ The impending catastrophic loss of biodiversity is no longer just a problem for ecologists and conservationists. Biodiversity loss and illegal wildlife trafficking have become threats to public health and food security as well. Public health and food security are under increasing threat from emerging diseases, with over 200 currently known zoonotic diseases, and 60 animal diseases that are listed by USDA as notifiable emergency situations.^2^ The World Bank estimates zoonotic diseases cause over $20 billion in direct losses to production animals and over $200 billion in indirect losses to related economies.^3,4^

Approximately three fourths of human emerging infectious diseases are caused by zoonotic pathogens, and 60% are shared with wildlife.^5^ While vectors and reservoirs for zoonotic diseases are diverse and complex, key underlying drivers for the increasing rate of spillover are habitat fragmentation, wildlife trafficking, bushmeat, climate change, and human encroachment on natural habitats.^6^ Fragmentation of forests and other habitats increases the interface between humans and wildlife, and the novel diseases those wildlife harbor. These intersections are an impetus for spillover^7^ – the larger this interface becomes, the more opportunities there are for a spillover event. Having lost approximately 10% of existing forest coverage since 1990, with losses disproportionately occurring in biodiversity hotspots,^8^ and when coupled with the rearrangement and distribution of ecological communities globally due to climate change, we may have irreversibly crossed a tipping point for spillover. While the tipping point, first characterized in containment efforts (i.e. *crush the curve* strategies) for SARS-CoV-2,^9^ typically refers to a critical number of infections in an outbreak wherein disease spread can no longer be contained and will grow exponentially, the emergence rate of all new zoonoses driven by climate change and biodiversity loss is following a strikingly similar pattern.^10^ In other words, if our climate and ecosystems degrade beyond this point, we may not be able to stop the increasing rate of emerging diseases and combating these threats to sensitive ecosystems, public health, and food security will become substantially more challenging.

Genetic testing is used to detect illegally trafficked wildlife,^11^ prevent the introduction of invasive species and pathogens,^12^ and monitor disease spread or outbreaks that can devastate the health of ecosystems, agricultural systems, and our communities alike.^13^ However, current models and platforms for forensics and diagnostics provide only a limited menu of available tests and are heavily restricted to centralized laboratories, far from where the threats to biodiversity are occurring and emerging. New tools, especially diagnostics,^14^ and novel approaches to biosurveillance, are imminently needed to protect our ecosystems and communities. Furthermore, these tools must prepare us for new unknown threats, while enabling early detection strategies that will minimize loss down the road.

To address this “neglected diagnostics” challenge our team at Conservation X Labs has developed a platform that can put the power of genetics into the field and into the hands of those at the front lines of these issues. This platform is composed of two key innovations that are presented here (Figure 1). The first is a handheld, battery powered tool that enables genetic tests to be performed by non-technical users at the point of contact. The Nucleic Acid Barcode Identification Tool (NABIT) provides a vehicle to move genetic testing outside of the laboratory and reduce the barriers to accessible and routine genetic testing anywhere, by anyone, at any time. However, a tool for decentralized wildlife genetics and biosurveillance is only as useful as the menu of readily available tests that can be performed to identify species, pests, and pathogen targets. To address this, our second innovation, the Molecular Development Kit (MDK), provides an avenue to resolve the limitations of available molecular tests by making the NABIT open to third-party assay developers in the global molecular diagnostic community who have laboratory assays they want to move into the field or desire to develop novel point-of-contact, field-based assays where early detection allows for greater impact and efficacy. The MDK is a genetic assay equivalent of software development kits (SDKs) which allows for the creation or translation of software applications for a specific platform, such as smart phones or video game consoles. In this case, our applications are molecular genetic ones. Like an SDK, our MDK is composed of documentation, software tools, and NABIT hardware that instruct and accelerate the other genetic assay developers to create shelf-stable tests for the NABIT that can be deployed into the field without a cold-chain. Herein, we demonstrate the performance of the NABIT and introduce the MDK.

**Figure 1:**
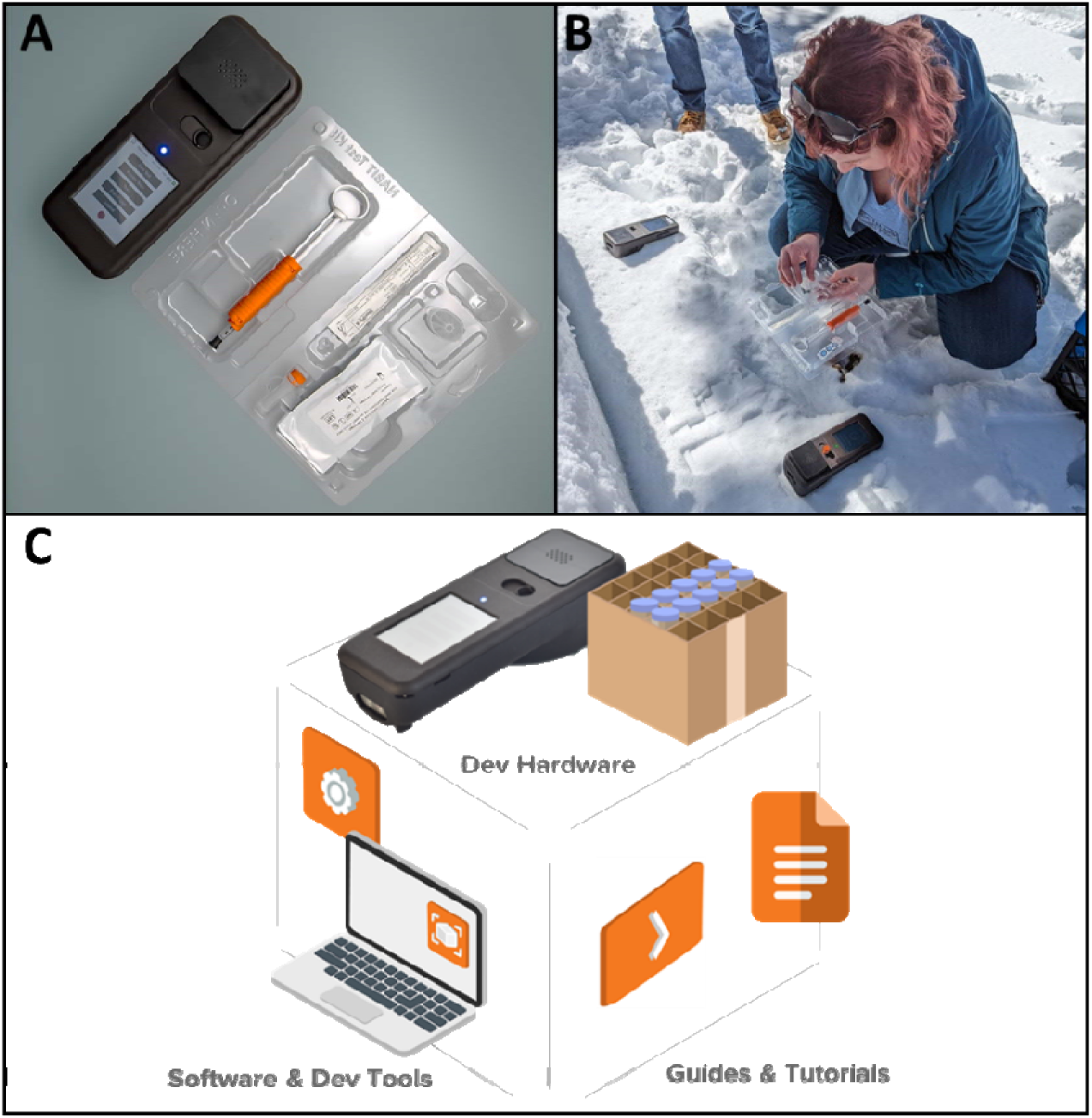
Decentralizing molecular testing for wildlife genetics and biosurveillance. The NABIT (A) provides a vehicle to enable point-of-contact genetic testing (B), and the MDK (C) provides a toolkit for the translation of laboratory assays or development of novel assays for field deployment.

## 2. Materials and Methods

### 2.1. Assembly of NABITs and Cartridges

The NABIT platform includes the durable instrument and test kits that contain cartridges specific to the target species or genetic sequence, which are inserted into the instrument for analysis.

NABIT instruments are manufactured using typical assembly techniques with custom and off-the-shelf componentry. The four major sub-assemblies of the NABIT are the 1) base assembly - consisting of the housing base, battery, and barcode scanner, 2) top assembly – consisting of the top housing, touch screen, and main electronic board, 3) core assembly (Figure 2) – containing the photodetector board, light emission channels, reaction heater, and lysis heater, and 4) door assembly – containing the excitation LED board, excitation optical channels (LED Guide), and a spring loaded compression mechanism to seal the reaction wells in the cartridge when the door is engaged.

**Figure 2:**
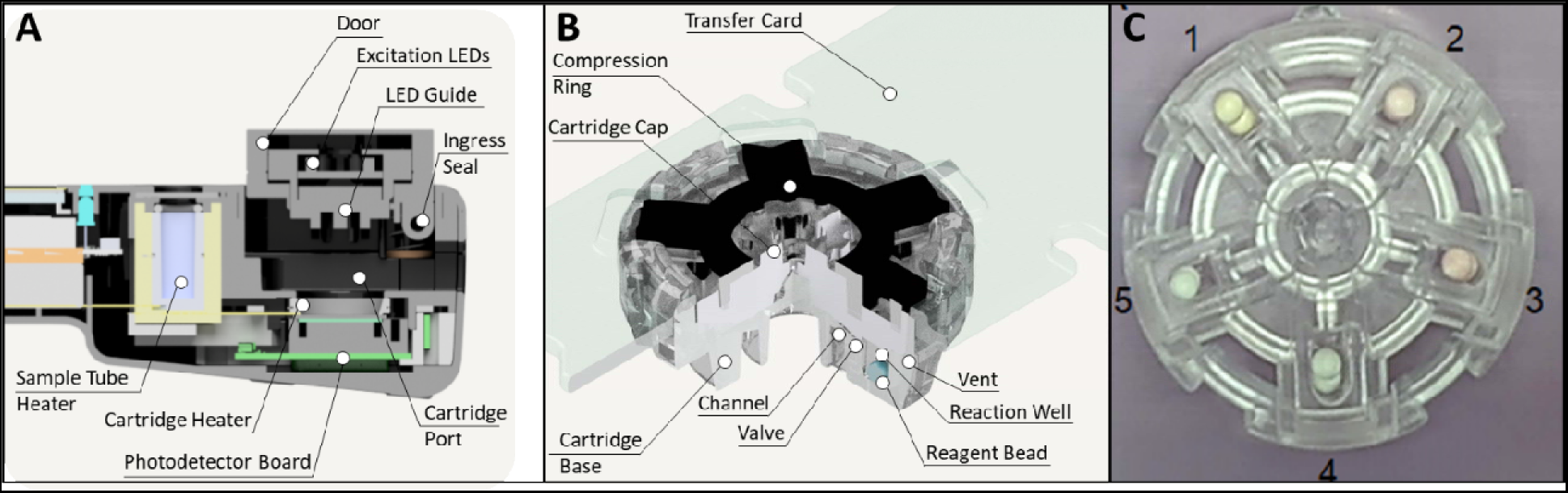
The NABIT enables sample to result tests with 5-well multiplex cartridges. The NABIT (A) provides a port to lyse samples in a collection tube and a reaction chamber that can heat cartridges and monitor a fluorescent reaction in each cartridge well. Assay beads are housed in cartridge wells (B-C). A channel is formed by geometry at the interface of the base and cap, enabling input sample to be distributed into each of the cartridge wells. A valve also formed at this interface seals the wells when the cartridge is compressed by the door assembly of the NABIT.

NABIT cartridges are composed of four components 1) a transparent, rigid base composed of cyclic olefin copolymer, 2) a clear flexible silicone rubber cap, 3) a rigid compression ring composed of black acetyl butyl styrene (ABS)/polycarbonate (PC), and 4) a transfer card composed of a black ABS. Each cartridge provides five wells that can house a specific assay for multiplex reactions.

### 2.2. Engineering Verification Tests

Following the assembly, we performed NABIT verification tests to characterize the key performance parameters of the instrument. These tests focused on measuring the thermal range and stability of the lysis heater and cartridge heater and the optical range of the photodetection system.

#### 2.2.1. Lysis Heater

To characterize the performance of the lysis heater we fed a single thermocouple (Omega #5TC-TT-TI-36-1M) through a 2 mL centrifuge tube cap (Thermofisher #3471TOS) and threaded onto a 2 mL centrifuge tube (Thermofisher #3490S) containing 280 µL of water. This thermocouple embedded tube is then placed into the lysis chamber in the NABIT to monitor the in-tube temperature while the lysis heater was activated. The thermocouple signal was recorded using DAQami data acquisition software (Digilent #6069-390-000).

#### 2.2.2. Cartridge Heater

We characterized the performance of the cartridge heater in a similar manner as the lysis heater by creating a thermocouple embedded cartridge with thermocouples placed central in the cartridge wells and embedded in adhesive. This thermocouple embedded cartridge is placed in the reaction chamber of the NABIT to monitor the temperature in each well of the cartridge while heating.

#### 2.2.3. Optical System

We interrogated the range of the optical system using cartridges that emit a range of fluorescence intensities with a serial dilution of fluorescein sodium ranging from 2 mM – 100 µM. For each cartridge, the reaction chamber is heated for 10 minutes allowing for 60 readings of each well to be taken by the NABIT.

### 2.3. Test Kit Implementation of Workflow

To enable a streamlined sample-to-result process with no measurement steps required by the user, we developed a test kit and custom transfer syringe around the workflow described in section 2.6. A holding tray houses the swab, lysis tube, custom transfer syringe, and detection cartridge. This tray also has a built-in guide flap to facilitate loading of the cartridge.

The custom transfer syringe enables users to add the diluent buffer to the lysed sample, and then withdraw the combined volume of 155 µL to be transferred to the cartridge without any measurements. This action is performed through a combination of cantilever stops built into a custom handle and finger loop that connect to a 1mL syringe (BD #309659). We make these parts from an injection molded ABS (Lustran 348). With this format the transfer syringe can input 1000 µL of diluent buffer into the lysis tube and when the plunger is withdrawn the cantilever restricts movement to ensure 155 µL is pulled within the syringe for transfer into the cartridge. We have also designed a skirt into the handle of the transfer syringe to prevent over insertion or overfilling of the lysis tube.

### 2.4. Software and Detection Algorithm

We developed an algorithm to automatically interpret the result of the amplification test for the user. In order to be adaptable to the variety of use cases the NABIT will be intended to address, we designed this algorithm to be specific and tunable enough to provide reliable results to specific assays as well as adaptable and adjustable to accommodate possible variances in behavior of different assays and sample types. The algorithm effectively detects signal in incremental step windows that identify positive reactions that have occurred in well channels (Figure 3). These step windows are defined by looking at the overall fluorescence signal derivative (dV_diff_/t), with defined signal derivative threshold (rate_Th_), signal steps would be labeled in tuple of start index and end index. Then, the width of a step, i.e., the duration of signal increase above derivative threshold (T_s_), is calculated by subtracting starting index from end index; the signal increase (V_diff_) is calculated from the integration of derivative within the labeled steps. Hence, the average signal increase rate (avgRate = V_diff_/T_s_) can be obtained. Within these step windows, we consider a set of threshold parameters (rate_Th_, width_LB_ and avgRate_Th_) to ensure the signal increase captured is from a true assay reaction. These variables look at the derivative of the reaction curve to filter potential false positives caused by an impulse to reaction signal, such as that caused by a mechanical shock or high-intensity signal interference occurring during the reaction, or signal drift that may occur during a reaction.

**Figure 3:**
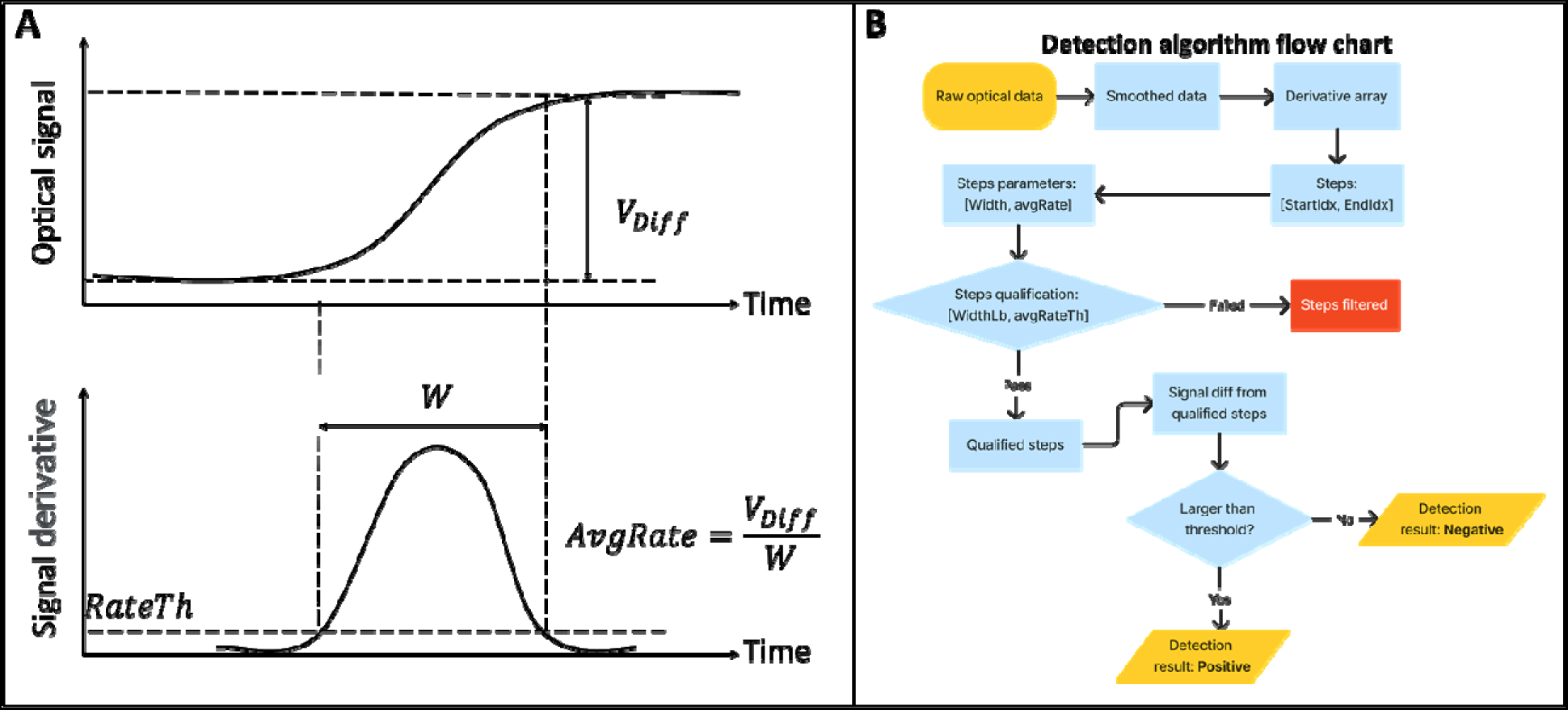
NABIT detection algorithm monitors fluorescent signal of reactions to provide automatic result interpretations and enable early call detection. The NABIT detection algorithms monitors for a sufficient rise in signal (V_diff_) to determine a positive test and analyses the derivative of the incoming signal to ensure signal rise occurs within a specific window (W) that is consistent with amplification behavior rather than in impulse or signal drift during the reaction (A). To call a result, the process performed by the algorithm is shown in a flowchart (B), where in the algorithm first performs this analysis on the behavior of the derivative and then checks that the signal threshold has been crossed to call a positive.

### 2.5. Lyophilized Assay Production

We used three reverse-transcriptase LAMP (RT-LAMP) assays to demonstrate the ability of the NABIT and cartridge to detect five reactions from a single sample. For the layout shown in Figure 2, Well 1 was loaded with an endogenous positive control that targets the human bacteriophage MS2 and contains a known concentration of RNA to monitor cartridge performance and inhibition. We loaded wells 2 and 3 with an assay that targets the nucleocapsid phosphoprotein (N) gene of the SARS-CoV-2 genome while the assay in wells 4 and 5 targets the membrane glycoprotein (M) gene in the SARS-CoV-2 genome. We describe the RT-LAMP primers in Table I.

We designed all assays for a 20 µL total volume. The assays are composed of 1X Isothermal Buffer (NEB #B0537), 6 mM Magnesium Sulfate (MgSO4 – NEB #B1003), 1.4 mM deoxynucleotide (dNTP) mix (NEB #N0447), 0.32 U/µL glycerol-free WarmStart Bst 2.0 (NEB #M0402B-BM), 0.3 U/µL glycerol-free WarmStart RTx (NEB #M0439B-HC1), 1X Chai Green (Chai Biosciences #R01200), 1X Excipient (Argonaut Manufacturing Services), and 1X primer mix (0.2 µM F3/B3, 0.8 µM LoopF/B, and 1.6 µM FIP/BIP). For the positive control, we included an MS2 RNA template of 5 x 10^6^ copies/µL (Sigma-Aldrich #10165948001) and 25 ppm of Orange G food dye (Sigma-Aldrich #O7252). The N gene assay included 25 ppm of Allura red food dye (Sigma-Aldrich #458848) so we may easily distinguish all three assay types from one another in bead form (orange, pink and uncolored). The total critical reagent volume was 10.183 µL, 10.133 µL and 10.033 µL for the positive control, N gene and M gene respectively.

Only the critical reagent volume is formulated for lyophilization since the sample is used to resuspend the LAMP reagents up to the final 20 µL reaction volume. We mixed all assays pursuant to their formulation, which we then dispensed as roughly 5 µL droplets into liquid nitrogen and lyophilized according to the manufacturer’s cycling and drying process (Argonaut Manufacturing Services – Carlsbad, CA). We split the critical reagent volume across two droplets so that the width of the lyophilized spheres (beads) would fit into the cartridge wells.

We manually dispense two beads into each well, either into five 0.2 mL optically clear qPCR tubes for evaluation on a Quantstudio 5 thermocycler (ThermoFisher), or directly into our cartridges using a tweezer-vac with a 2.38 mm vacuum cup (Virtual Industries #TV-1000-110 and V8903-D-S) in an isolation glove box (Cleateach #2100-2-E) maintained at <10% RH according to the cartridge layout in Figure 2.

To ensure the stability of bead performance over time, we tested the performance of assembled cartridges (N=54) that had been stored at room temperature (20°C) for 17 months after bead production and cartridge assembly.

### 2.6. Molecular performance tests and controls

We conducted quality control checks of bead quality prior to cartridge loading with synthetic template standards of SARS-CoV-2 RNA (TWIST #102024) diluted in UltraPure™ DNase/RNase-free distilled water (UP_d_H_2_O - Invitrogen #10977015). To evaluate the performance of the NABIT reaction system, we diluted samples of gamma-irradiated SARS-CoV-2 virus isolate USA-WA1/2020 (BEI #NR-52287) in human nasal wash (Lee Biosolutions #991-26-P) which were used as reference samples. No template controls (NTC’s) utilized UP_d_H_2_O or human nasal wash only.

For synthetic template standards, we created a 1:10 dilution series ranging from 100,000 to 100 copies/µL (input [C]). To mimic a sample lysate, a custom lysis buffer consisting of 25 mM UltraPure^TM^ Tris-HCl (Invitrogen #15568025), 25% QuickExtract^TM^ RNA (Biosearch Technologies #SS000880-D2) and UltraPure^TM^ DNase/RNase free distilled water (Invitrogen #10977015) was heated to 95°C followed by the addition of a custom diluent buffer consisting of 25 mM UltraPure^TM^ Tris-HCl (Invitrogen 15568025), 1X RNAsecure^TM^ RNase Inactivation Reagent (Invitrogen #AM7005) and UP_d_H_2_O (Invitrogen #10977015) (Invitrogen #10977015) at a ratio of 1:5. 18 µL of the contrived sample lysate was combined with 2 µL of diluted synthetic standards to resuspend each assay type in duplicate at final RNA concentrations of 10,000 to 10 copies/µL. These standards were incubated on a QuantStudio^TM^ 5 Real-Time PCR thermocycler (Applied Biosystems #A47327) at an annealing temperature of 65°C with fluorescence measured every 60 seconds for 45 minutes (min), followed by a melt curve analysis.

Range finding and performance testing was conducted by the Atlanta Center for Microsystems Engineered POC Technologies at Emory University (ACME-POST - Atlanta, GA) under the auspices of the Rapid Acceleration of Diagnostics (RADx) program. A sample-to-result workflow was used with gamma-irradiated virus standards by loading 50 µL each of a dilution series ranging from 7,650 to 12 NDU/µL onto a polyester swab (Puritan #25-806). This swab was dipped into a 2 mL microcentrifuge tube with tethered O-ring cap (FisherScientific #3490 and 3471TOS) containing 400 µL of the custom lysis buffer for 10 seconds and then removed. The tube was heated to 95°C for 5 min in the NABIT lysis chamber followed by addition of 1 mL of diluent buffer using the transfer syringe in 2.3. Finally, 155 µL +/- 5 µL of the prepared standard was loaded into the cartridge at final concentrations ranging from 29.77 to 0.47 NDU/µL. The cartridge was inserted into that NABIT and heated to 65°C for 30 min followed by result interpretation by the algorithm.

For long-term storage and stability tests, we performed the sample-to-result workflow described above with two gamma-irradiated virus standards at 7,650 NDU/µL or 1000 NDU/µL, or an NTC with human nasal wash.

## 3. Results and discussion

The NABIT provides a hand-held, battery powered platform that can detect genetic targets with an onboard sample preparation chamber to enable complete samples-to-results tests (Figure 4) of many sample types without additional equipment. The integrated touchscreen provides an accessible user interface. The accompanying NABIT cartridge performs five simultaneous reactions, enabling multiplex targeting (for more sensitivity or specificity, different tests, or redundant reactions for increased reliability). The design of the NABIT cartridge in combination with the use of lyophilized assay beads provides for an agile manufacturing platform for new test layouts to be readily assembled when additional assays become available.

**Figure 4:**
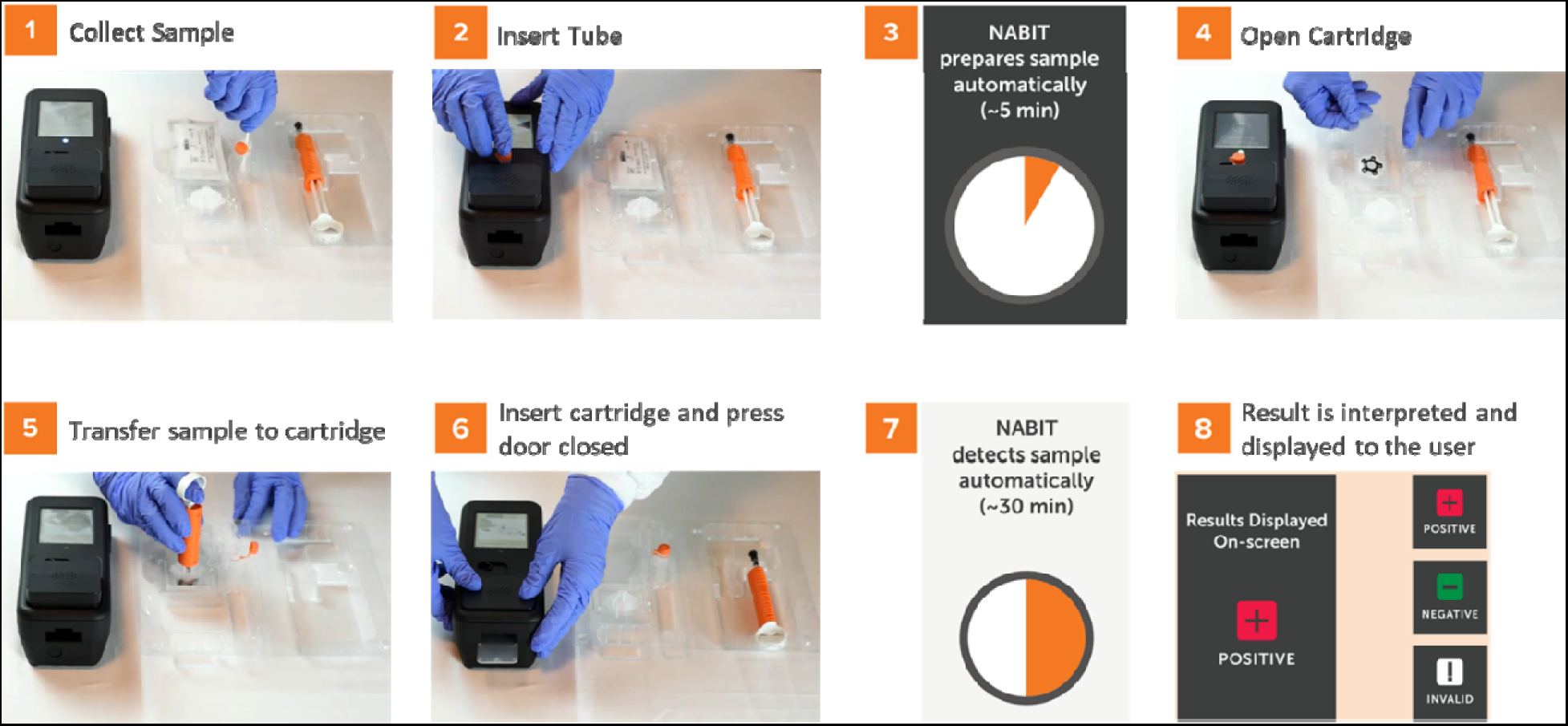
The NABIT and accompanying test kit provide a streamlined workflow to run genetic tests without precision measurements or techniques. A test is run on the NABIT first collecting and inserting the sample into the lysis tube (1) and inserting the tube into the NABIT (3) for thermal lysis (4). During lysis, the cartridge can be removed from its hermetically sealed pouch (4), upon which a fail-safe transfer syringe is used to add diluent and transfer the processed sample into the cartridge (5). The loaded cartridge is inserted into the NABIT (6) where the reaction is automatically performed (7) and result displayed to the user (8).

### 3.1. Sample preparation

The sample preparation chamber consists of a heatblock and heating element that supports a standard 2mL sample tube. The temperature is controlled by the NABIT software and monitored by a thermistor on the heatblock. The chamber can safely reach temperatures up to 100°C. In-tube temperature measurements show that the NABIT sample preparation chamber can reach and maintain 95°C within ± 2°C through the completion of the heating cycle (Figure 5). Furthermore, the ramp time from room temperature to 95°C was determined to be 3 minutes and running a 95°C lysis step for 5 minutes was measured to consume 250 mAh. The ability to reach 95°C and stably maintain this temperature demonstrates the ability of the NABIT to enable preparation of sample types that require inactivation^15^ at high temperatures and covers the range of temperatures used in thermolysis.^16,17^

**Figure 5:**
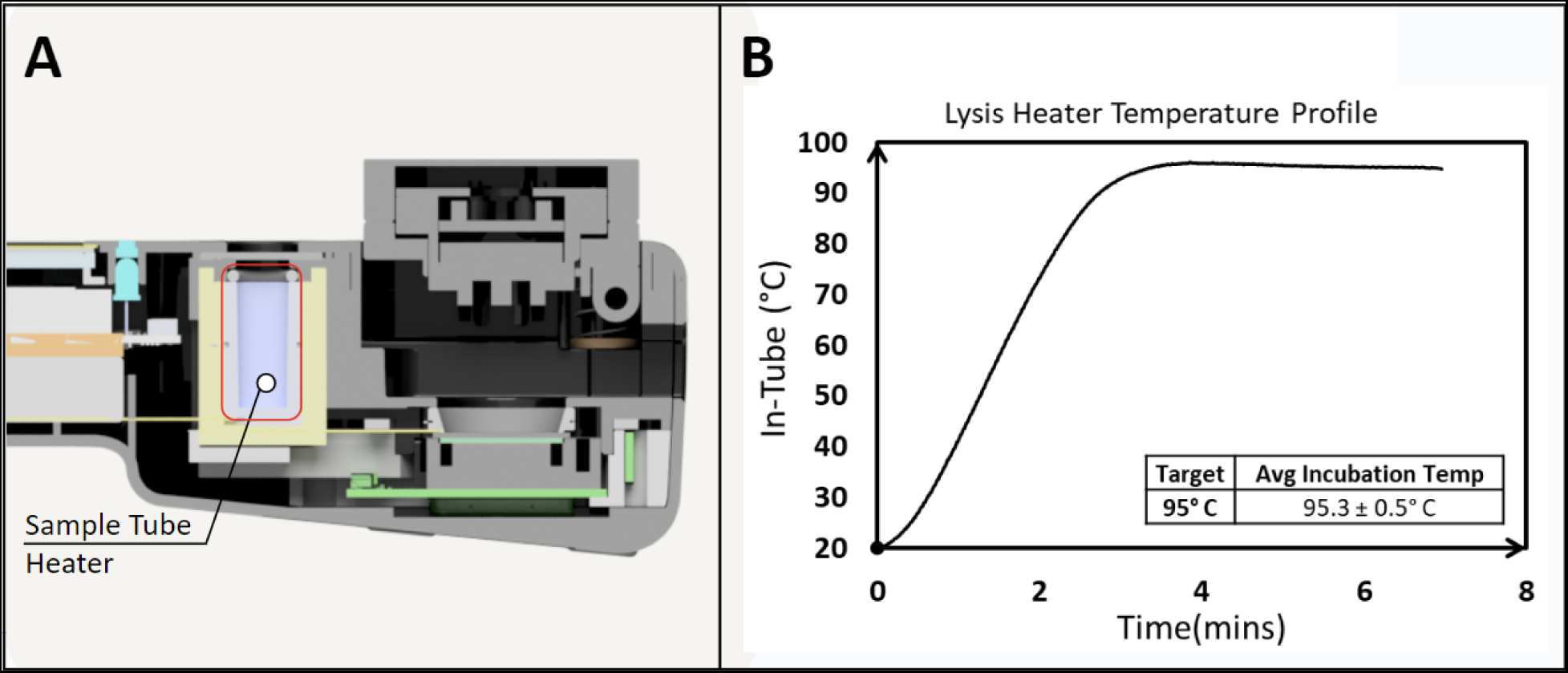
On-board NABIT lysis heater provides quick ramp up to high temperatures for thermal sample processing. The sample tube heater (A) demonstrates a ramp time up of 180 seconds to maintain a lysis temperature of 95.3°± 0.5°C.

### 3.2. Reaction-detection chamber

The temperature tests on the reaction-detection chamber were measured using the thermocouple-embedded cartridge as described in section 2.2.2. Results demonstrate that the cartridge heater could maintain temperatures up to 70°C with a well-to-well variance of ± 1.0°C (Figure 6). The reaction-detection chamber takes less than six minutes for each well to reach the target reaction temperature. The detection system of the NABIT could detect concentrations of Fluorescein down to 2 µM and as high as 10 µM, sufficient to cover the emission range of a LAMP reaction with an intercalating dye. Together, the reaction chamber exhibits the required temperature range and stability to reliably drive isothermal amplification tests and monitor fluorescent emission of indicator dyes throughout the course of the reaction. This ability, combined with an automated detection algorithm also enables early call functionality to be implemented on the NABIT.

**Figure 6:**
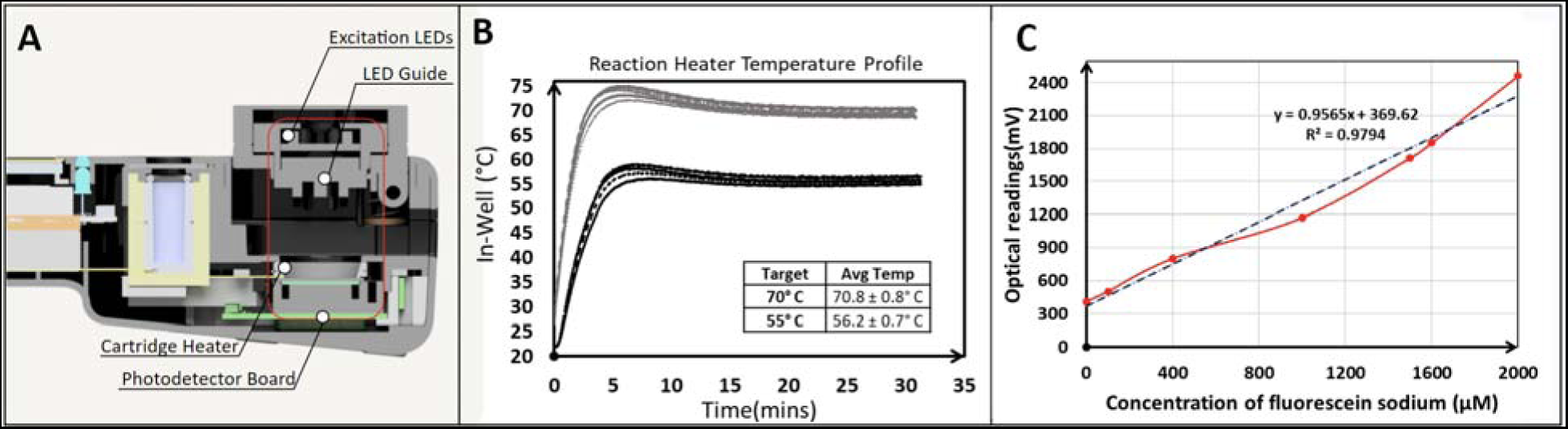
The NABIT reaction-detection chamber provides stable thermal profiles and optical sensitivity to drive and monitor amplification reactions. The NABIT reaction-detection chamber (A) provides ramp times less than 6 minutes for reaction temperatures ranges from 55 to 75°C and maintains target temperatures to ± 1.0°C (B). The optical sensors detect fluorescein sodium with a linear trend (R^2^ = 0.98) across a concentration of 2 to 10 µM (C).

### 3.3. Test Performance

Quality control tests using TWIST synthetic RNA confirmed the expected performance of lyophilized beads. The positive control started amplification at an average of 11.4 min (SD 0.18) across 32 replicates. The N gene and M gene assays showed consistent amplification (at least 3 of 4 replicates) of concentrations between 10,000 to 10 NDU/µL between 11-20 minutes (SD 1.31 and 1.11 overall), and no amplification in the NTC replicates.

Performance testing at ACME-POCT required at least 3 positive tests to confirm a positive result for each standard concentration. The results demonstrated that the NABIT could detect template targets down to a concentration of 0.93 NDU/µL (Figure 7), the equivalent of about 18.6 NDU in a reaction well, a value that is approaching the theoretical limit of 10 copies required for LAMP reactions.^18^ The NABIT can detect lower concentrations, as shown by the 0.47 NDU/µL standard, but not as consistently. A false positive was reported in a negative sample that was attributed to the inherent contamination risk involved with diagnostic sampling. We will continue to characterize these results and adapt with changes to the sample workflow and detection algorithm to ensure a low rate of false positives.

**Figure 7:**
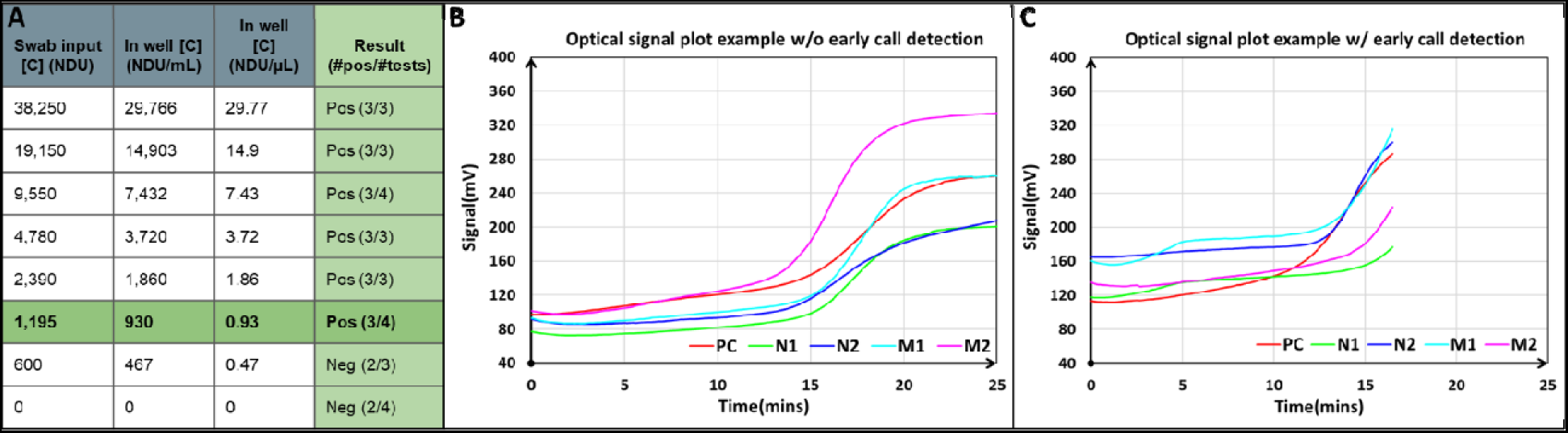
The NABIT can provide sensitive, low copy number detection of input samples. Rangefinding limit of detection tests performed by ACME-POCT at Emory University demonstrated detection of input sample down to 0.93 NDU/µL (A). Fluorescent curves on the NABIT show the monitoring and early call capability of a positive result (B).

Stability tests produced 9 out of 9 positive (100%) results from the 7,650 NDU/µL standard, 27 out of 28 (96.4%) positive results from the 1000 NDU/µL standard, and 17 out of 17 negative results from the NTC nasal wash. One test produced an invalid result due to user error. This demonstrates that the lyophilized beads can retain performance after 17 months when stored at room temperature and hermetically sealed.

### 3.4. Creating New Assays with the Molecular Development Kit (MDK)

Having demonstrated the performance of the NABIT and long-term stability lyophilized assays, we can reproduce the process of developing additional field-ready test kits (Figure 8) for the NABIT for a wide variety of sample types and targets in wildlife genetics and biosurveillance. The goal of the MDK is to standardize that development of new assays, or transfer of existing laboratory-based assays, into field-ready test kits for the NABIT and make this process accessible and as simple as possible (and even automated for some steps) for other scientists to also reproduce and create their own field-ready tests.

**Figure 8:**
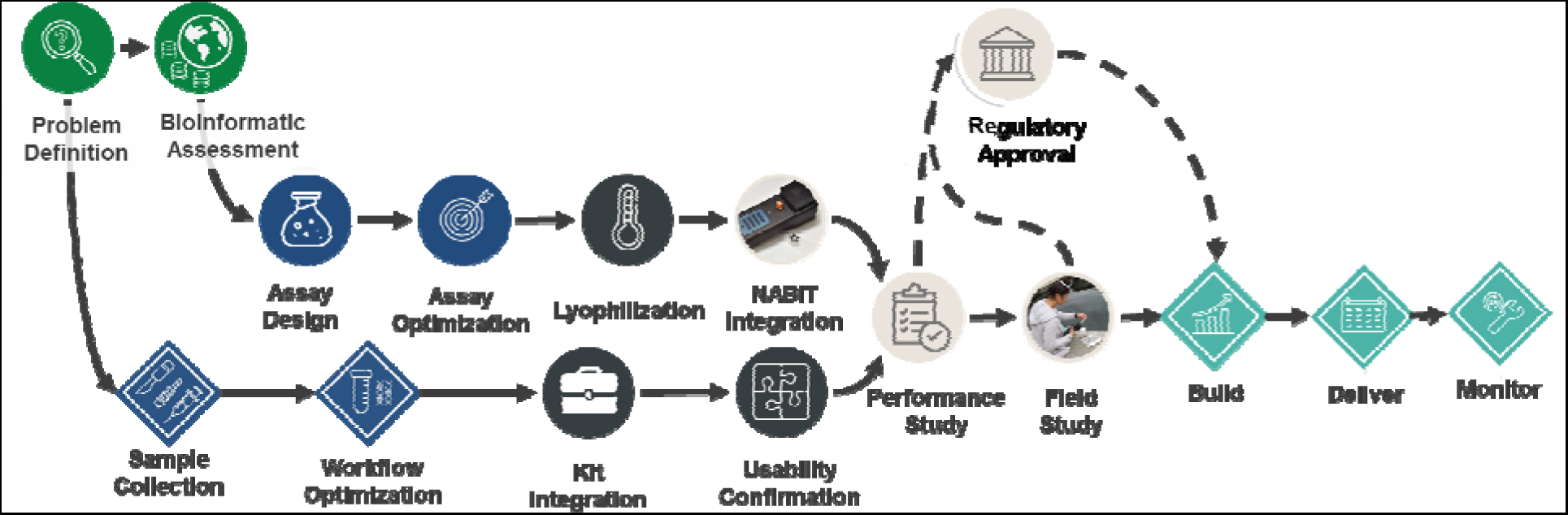
An overview of the development and deployment process of field-ready test kits with the NABIT.

As part of the MDK, starter packs will provide a sample of available cartridges, kit components, and recommended buffers to facilitate this process for third-party developers, along with the NABIT hardware. We will also provide supporting software tools to guide developers through the process of determining and designing assay targets and setting parameters on NABIT hardware, including customizing the animated graphical user interface and training the detection algorithm on their specific assays (Figure 9). Tutorials and documentation will accompany each of these tools and provide a comprehensive instruction set and troubleshooting tips for each stage of development of a NABIT test kit.

**Figure 9:**
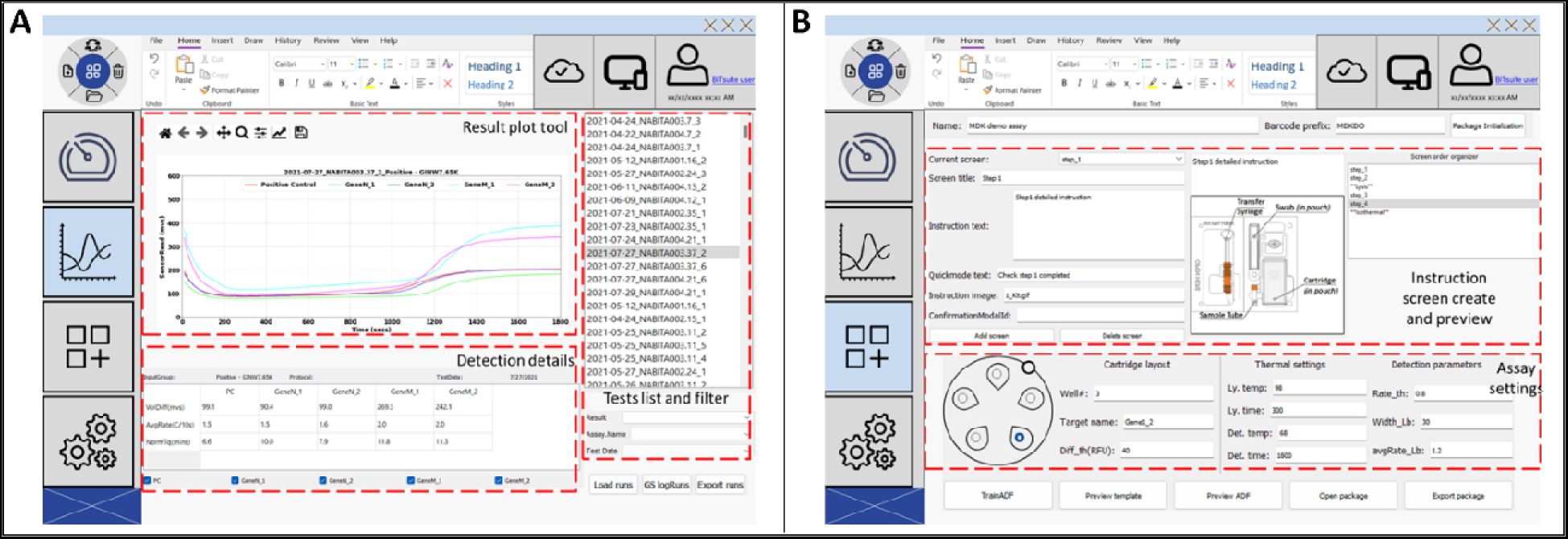
Desktop MDK software tool enables developers to program NABIT workflows. Screen shots of prototype software tool that allows users to optimize the NABIT parameters and train the detection algorithm for their assays (A), and program the graphical user interface that end users will see for their specific tests (B).

A key feature of the NABIT that enables the MDK and this model to address neglected diagnostics is a containerized software architecture (Figure 10) that has been developed for the NABIT to allow for facile updates of new assays, data interpretation, and features to be implemented on the system to meet the needs of a variety of users and developers. This model also enables the GUI to be run through a kiosk mode, providing additional security from unintended or unauthorized use of the device and tests.

**Figure 10:**
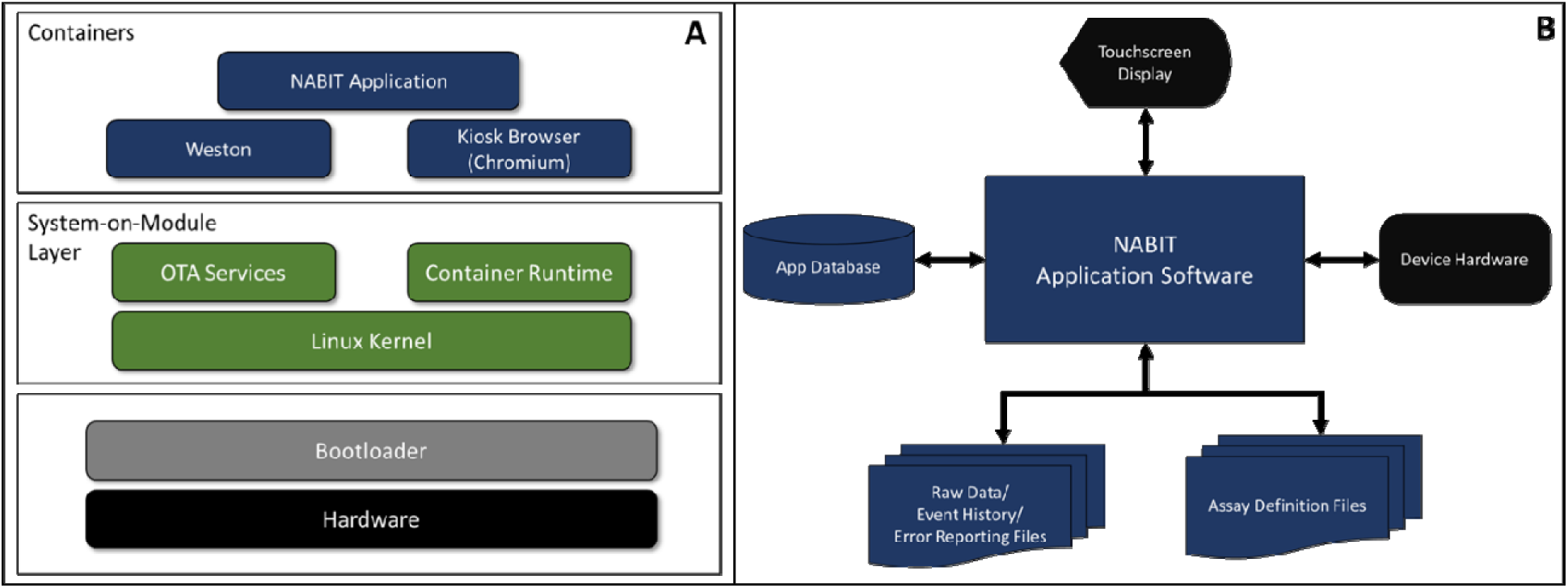
NABIT software architecture supports extensive library of unique test workflows. The NABIT software stack is built off of a Linux Kernel to enable a containerized software architecture (A). The containers for the NABIT Application Software call data from an Assay Definition File, that is uniquely created for each test during the assay development process, to operate NABIT hardware components and generate report files containing the data from each test and any errors detected or metadata input during the performance of the test (B).

## 4. Discussion

Overall these results lay out the potential of the NABIT to address critical needs for on-site testing in applications such as one health for wildlife and veterinary disease, pandemic surveillance, and combating wildlife crime in trade and traceability. The modularity of the NABIT software architecture and format of test kits and cartridges provides for facile adaptation and implementation of new assays and test layouts into a test kit.

However, the NABIT by itself is not a complete solution to the neglected diagnostic challenge. To effectively address this challenge, a broad menu of ready-made and effective diagnostic tests is required to meet the plethora of species and disease targets in need. Traditional diagnostics business models have failed to address this immense diversity in diagnostic demand because manufacturers focus on producing a small menu of high-demand tests where they can maximize profits. These closed-door and market-response approaches for diagnostics may not be the best suited model for critical diagnostic needs such as diseases of the developing world, including tuberculosis and malaria, because most patients suffering from these diseases have poor paying capacity^19^. This paying capacity is even further limited for applications in conservation and for non-human diagnostics of emerging pathogens that have yet to infect many people. Selecting for the highest return on investment has resulted in intense competition for a small number of diseases and use settings, leaving a long tail of potential diagnostic tools uncompleted, and with it, undermining the possibility of addressing biodiversity loss or pests and pathogens that affect wildlife, domestic livestock, agricultural plants, and ultimately, undermine human security and planetary health.

The purpose of the MDK is to enable a new model for making effective diagnostics more accessible, more available, and more diverse. We realize that our team will never solve this challenge if we focus on continually developing a sequence of new tests for the NABIT ourselves. By opening NABIT development tools that allow the global community to develop or translate their own tests for field use on the NABIT, we can harness the collective talent and efforts of many scientists and experts working on the same challenge in their own silos. This approach was inspired by smart phones (e.g. the Apple App Store) and the video game industry, wherein publishers like Sony or Microsoft host consoles for which many third-party developers can design unique titles.

The video game industry would not be what it is today if a community of commercial game development companies, independent game developers, and even accessory hardware designers and producers had not grown alongside the console manufacturers to support gamers and enhance their experience. By using this wildly successful model to solve neglected diagnostics, we can similarly grow a development community for diagnostics tests and services. While our MDK platform will serve as a key toolkit to inspire and empower others to develop tests, we will also provide a comprehensive set of options to engage our development community and support a wide range of needs and capabilities to usher along new tests into commercial availability and global accessibility. From this foundation, we propose creating a larger global coalition of partners around the MDK. Our goal is to not only to develop new diagnostics, but to build and motivate a community, harness the power of solvers across industries that continue to address the neglected diagnostics challenge, and ensure the innovations find markets that can help them scale. This vision can only be achieved through a strong, interconnected, and collaborative network across academic institutions, multilateral institutions, governments, investors, corporations and non-profits.

## Author Contributions

Bunje PME, Dehgan A, Baisch DB, Winters M, Fang C, Fotouhi G, and Holmes HR conceived the ideas; Baisch DB, Winters M, Mercader J, Fang C, Fotouhi G, and Holmes HR developed the methodology; Winters M, Mercader J, Fang C, Fotouhi G collected the data; Baisch DB, Winters M, Mercader J, Fang C, Fotouhi G, and Holmes HR analyzed the data; Dehgan A and Holmes HR led the writing of the manuscript. All authors contributed critically to drafts and gave final approval for publication.

## Acknowledgements

The authors would like to acknowledge Fahim Farzadfard and Louis Kang for their assistance in primer design and the Atlanta Center for Microsystems Engineered Point-of-Care Technologies (ACME-POCT) for the assistance in assay validation. The authors would like to acknowledge the Gordon and Betty More Foundation, the Schmidt Family Foundation, the National Institutes of Health Rapid Acceleration of Diagnostics (RADx Next) Program, KKR Philanthropy, and the Center of Complex Interventions for their support of this work.

---

**Table S1:**
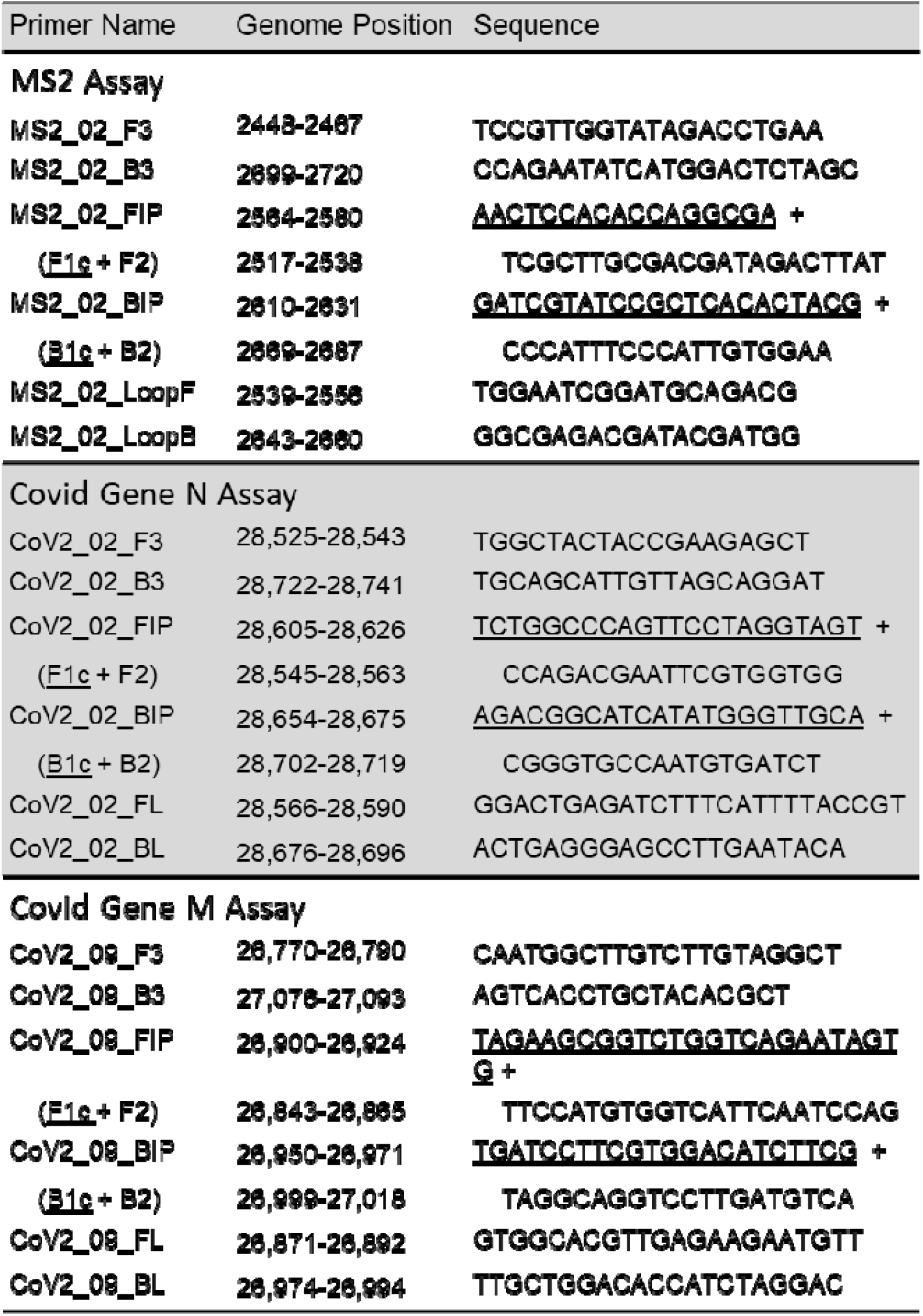
RT-LAMP primers and target positions.

