## Supplementary figures and images for "Decentralizing genetic testing for biodiversity monitoring and biosurveillance with the Nucleic Acid Barcode Identification Tool (NABIT) and Molecular Development Kit (MDK)"

### Supplementary Table 1

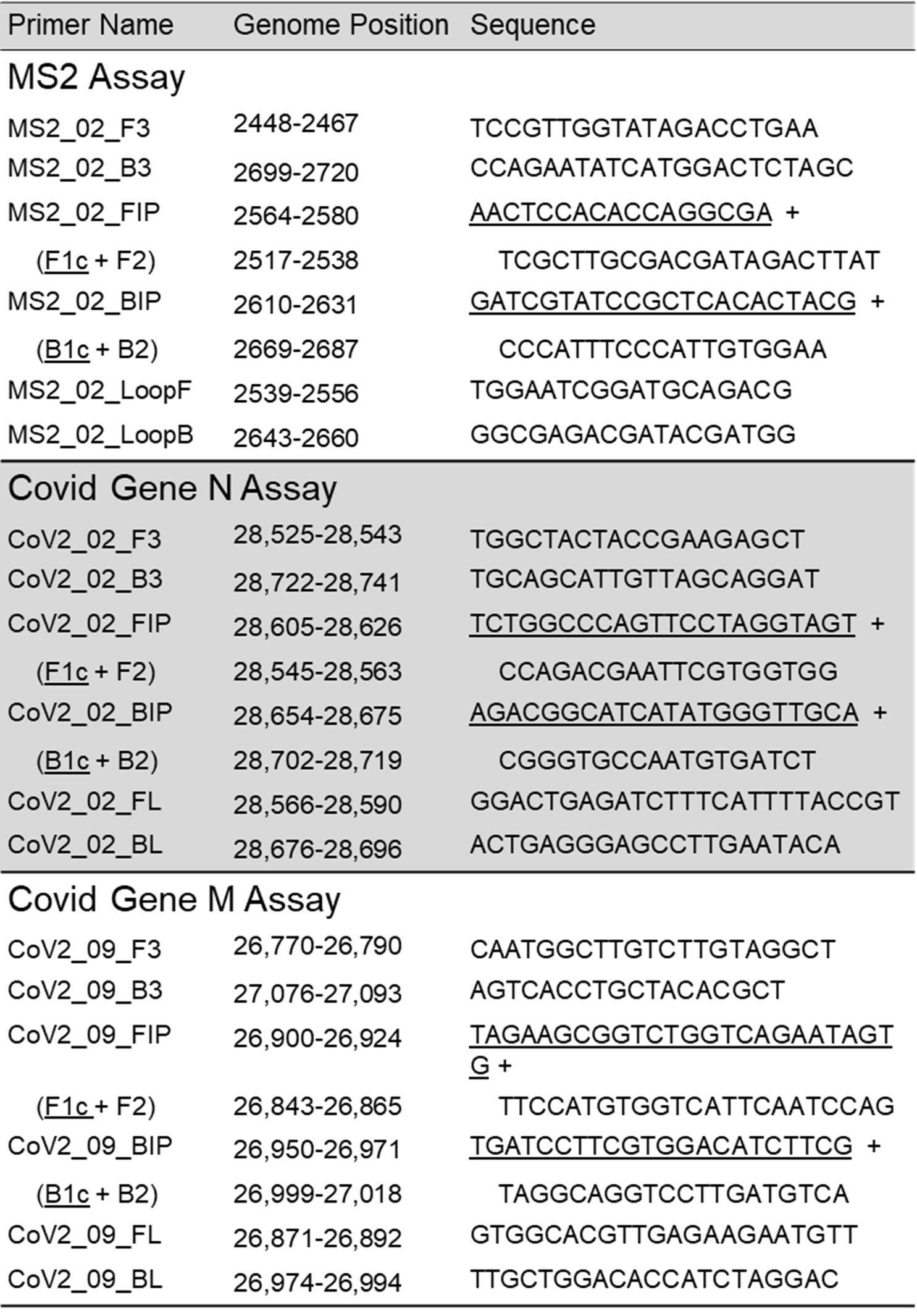
